# Establishing an RNA sensor with high sensitivity and dynamic range utilizing a signal amplifier platform

**DOI:** 10.1101/2025.09.17.675666

**Authors:** Ha Eun Lim, James Chappell, Laura Segatori

**Affiliations:** Department of Bioengineering, Rice University, Houston, TX, 77005, USA; Department of BioSciences, Rice University, Houston, TX, 77005, USA; Department of Chemical and Biomolecular Engineering, Rice University, Houston, TX, 77005, USA

**Keywords:** RADAR sensors, gene signal amplifier, mammalian synthetic biology, RNA sensors

## Abstract

Precise control of gene expression in a cell-state-specific manner is essential for effective therapeutic interventions in complex and dynamic disease microenvironments. Traditional targeting strategies that rely on surface markers or cell type-specific promoters often assume static cellular identities, limiting effectiveness in contexts such as cancer and inflammation, where cell states are highly heterogeneous and dynamic. RNA sensors, such as RADAR (RNA sensing using Adenosine Deaminases Acting on RNA), provide a modular, programmable, and non-integrating platform for classifying cell states. However, it is also characterized by low sensitivity and dynamic range, which limits its applications in detecting low-abundance transcripts. In this work, we integrate RADAR sensors with a signal amplification circuit to enhance sensitivity and dynamic range. We demonstrate that this combined RADAR-amplifier platform enables real-time monitoring of subtle changes in the abundance of endogenous transcripts under physiological conditions. Our results demonstrate the utility of this platform for fundamental biological studies and the development of precision therapeutic strategies.

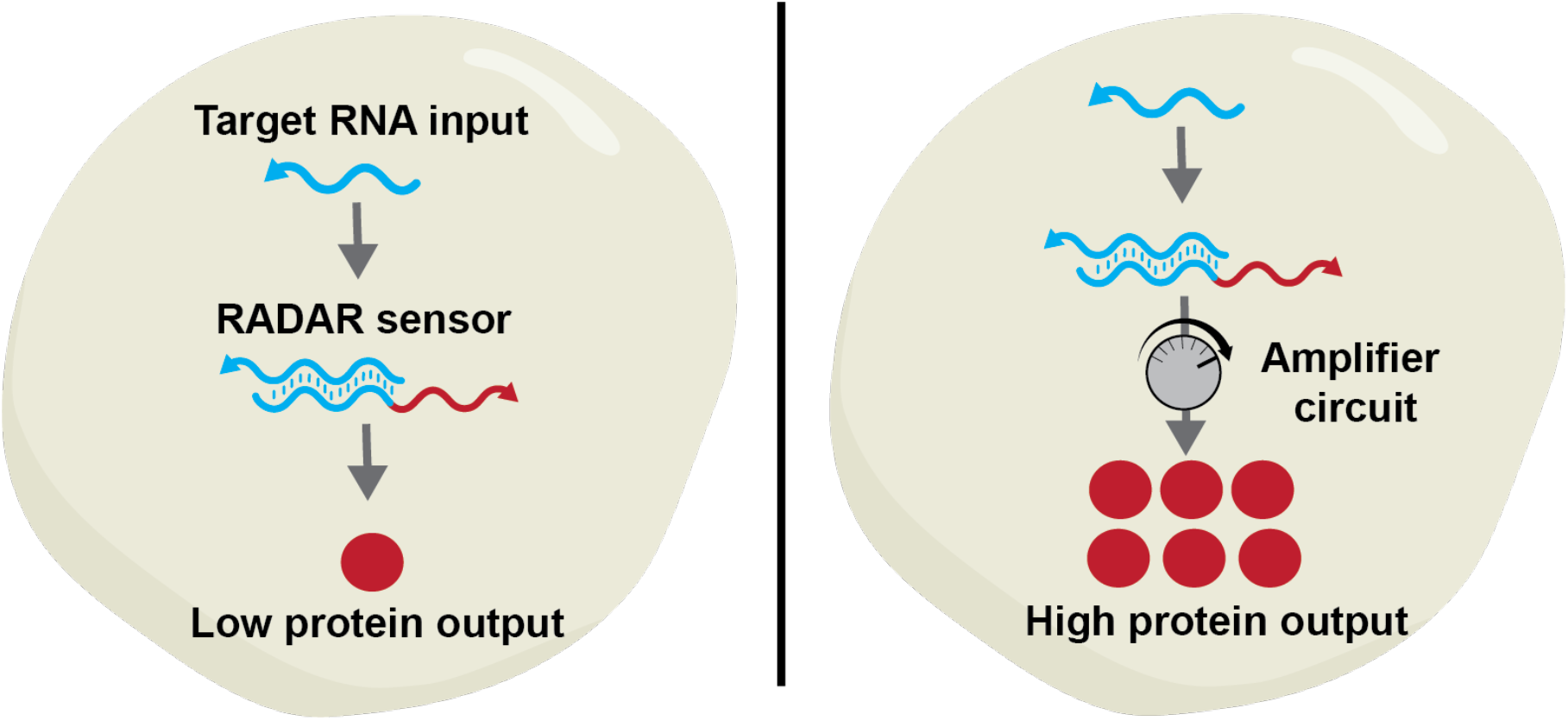

## Introduction

The ability to precisely regulate therapeutic payload expression in a cell-state-specific manner is a critical requirement for achieving targeted interventions. Unlike conventional strategies that broadly activate therapeutic programs across diverse cell populations, cell-state-specific systems are programmed to target cells exhibiting defined molecular signatures. Developing therapeutic strategies with high specificity is particularly crucial in complex tissue environments characterized by cellular heterogeneity and dynamic cellular states. Such complexity is a hallmark of many diseases, such as cancer and autoimmune disorders, as well as natural cellular processes, such as tissue regeneration.^1–3^ Solid tumors, for instance, comprise a heterogeneous mixture of cancer stem cells, differentiated tumor cells, stromal components, and infiltrating immune cells— each playing distinct roles in tumor progression, immune evasion, and therapeutic resistance.^4^ Similarly, autoimmune pathologies involve dysregulated immune responses driven by transient subsets of pro-inflammatory cells within microenvironments also comprising immune cells with protective roles.^5^ In these settings, therapies that lack cell-state specificity are inevitably plagued by low therapeutic efficacy and high side effects.

Cell-state classifiers programmed to detect and respond to specific molecular features that define a cell’s functional state offer a promising strategy for dynamic and selective targeting.^6^ Cell-state classifiers comprise a diverse set of tools designed to sense molecular features that define a cell’s current functional or pathological state, and in response, control downstream gene expression programs. By responding to endogenous molecular cues—such as protein expression profiles—rather than fixed cell-type markers, cell-state classifiers provide a framework for developing therapies that are both dynamic and specific to disease-relevant cell populations.

Current strategies for implementing cell state-specific targeting typically rely on engineered viral or non-viral vectors that recognize surface markers unique to specific cell types.^7–9^ Alternative targeting approaches utilize cell type-specific promoters or enhancers to drive selective transgene expression.^10–12^ While these approaches have enabled significant advances in cell type-specific delivery, they are often limited by critical constraints. Surface marker-based targeting requires detailed knowledge of unique, stably expressed cell-surface antigens, which may not always provide unique molecular signatures.^13^ The use of cell type-specific promoters or enhancers necessitates extensive screening and validation to ensure sufficient specificity and expression strength, making the development process labor-intensive and non-generalizable.^14^ Moreover, these strategies typically assume static cell identities and do not account for the dynamic and transitional nature of cellular states, which is characteristic of complex disease states such as cancer and inflammation.^14,15^ As a result, they may exhibit low effectiveness in diseases characterized by plasticity and heterogeneity, resulting in suboptimal targeting and therapeutic escape.

Recent advances in single-cell transcriptomics have provided powerful tools to define cell states by their unique RNA profiles,^16^ paving the way for a new class of cell-state classifiers based on RNA sensors that link RNA signatures to programmable gene expression outputs. RNA sensor-based classifiers offer several advantages over traditional approaches: they are highly modular, non-invasive since they do not require genomic integration (minimizing insertional mutagenesis risks),^17^ can be designed rapidly and *de novo* using well-defined base-pairing rules,^18^ and can recognize target RNA even in the presence of high mutational burden, a common feature of plastic environments such as tumors and inflamed tissues. As a result, RNA sensors represent a promising and versatile platform for cell-state classification and precise therapeutic targeting.

The toolkit of RNA sensors has expanded rapidly in recent years: toehold switches that control translation by conditionally exposing translation initiation sequences in the presence of complementary RNA sequences,^19–21^ split-ribozymes that control gene splicing,^22^ miRNA toehold switches that tune expression based on endogenous small RNA levels,^23^ and CRISPR-Cas9-based RNA sensors that actuate genome editing or transcriptional regulation in response to guide RNAs.^19,24^ More recently, a new class of RNA sensors has been developed, known as RADAR (RNA sensing using Adenosine Deaminases Acting on RNA) sensors.^25–27^ These sensors leverage the activity of endogenous or engineered ADAR enzymes, which catalyze site-specific adenosine-to-inosine editing in response to the presence of a target RNA. RADAR platforms are highly modular and programmable: sensors can be rapidly designed *de novo* to target virtually any RNA of interest, and the outputs—ranging from diagnostic fluorescent and luminescent reporters to therapeutic molecules such as caspases, recombinases, or small peptides—can be easily interchanged. These properties make RADAR sensors a promising tool for real-time cell state monitoring and the associated implementation of programmable actuations. However, a key limitation of RADAR sensors is their relatively low sensitivity for detecting low-abundance endogenous transcripts.^26,27^ While effective for sensing transcripts expressed at high levels from plasmids or synthetic constructs, their performance for monitoring endogenous RNAs with low or transient expression, which are often characteristic of cell states, remains suboptimal. This limited sensitivity is paralleled by a narrow dynamic range of the output signal, which hinders the ability of RADAR sensors to distinguish small but biologically relevant changes in transcript levels.

To address the limitations of RADAR sensors, we explore the use of a genetic circuit specifically designed to function as an output signal amplifier.^28^ Since improving sensitivity to small changes in RNA levels and expanding the output signal dynamic range of RADAR sensors are critical, we hypothesized that incorporating an output signal amplifier would improve the detection of subtle variations in the input signal and amplify the dynamic range. This strategy has been previously shown to enhance the performance of conventional reporter systems, which directly couple the expression of a diagnostic reporter to that of a target gene and are therefore constrained by the often low expression level of the target, resulting in correspondingly low reporter signal. A gene signal amplifier strategy, which links the expression of the target gene to that of an orthogonal genetic circuit designed to amplify the output signal through the integration of transcriptional and post-translational control elements, can potentially enhance the output signal’s dynamic range and improve the resolution of input signal variations. Integrating such signal amplification technology with the RADAR sensor to generate a RADAR that triggers a genetic circuit for output signal amplification is expected to enhance the RADAR’s sensitivity to changes in endogenous transcript levels, thus addressing the limited sensitivity and narrow dynamic range that currently restrict applications of this technology for real-time detection of cell states.

In this study, we systematically evaluated the performance of an integrated RADAR-amplifier platform. We first optimized the output dynamic range and compared the RADAR sensor integrated with the signal amplifier circuit to an uncoupled RADAR sensor. We then applied this integrated platform to detect changes in endogenous transcript levels under biologically relevant conditions. By enhancing both the input resolution and sensitivity of RADAR sensors, this work expands the applicability of RNA-based classifiers for the precise monitoring of cell states based on defined molecular signatures, with broad applications as cell classifiers in heterogeneous and dynamic environments.

## Results

### Engineering RNA sensors with optimized output signal dynamic range and sensitivity

To engineer RNA sensors with a low limit of detection and high dynamic range that enable monitoring small fluctuations in target RNA levels, we leveraged a previously developed RNA sensor, <u>R</u>NA sensing using <u>A</u>denosine <u>D</u>eaminases <u>A</u>cting on <u>R</u>NA (RADAR).^26,27^ RADAR sensors require two components for detection: the RNA sensor sequence and the ADAR enzyme, typically delivered into cells using two plasmids (Fig. 1a). The RADAR sensor is activated only upon interaction with the target RNA sequence, resulting in the production of a user-defined output protein. Specifically, the RADAR sensor sequence comprises three main elements: (1) a sequence encoding a reference protein that is expressed constitutively and independently of the RNA sensor activity, providing a reference to quantify the extent of plasmid delivery and expression level of the RNA sensor, (2) a sensor sequence that is reverse complementary to the target RNA and contains a premature stop codon (UAG) to block downstream translation, and (3) a sequence encoding the output protein, that is expressed only upon detection of the target RNA. The RADAR sensor plasmid is co-delivered with a plasmid encoding the ADAR enzyme, which in this study was designed to overexpress the p150 isoform of ADAR1, as it was expected to provide an improvement in RADAR activity as previously demonstrated.^26,27^ In the absence of the target RNA, the stop codon within the sensor RNA sequence causes premature termination, thereby preventing the translation of the output protein. The ADAR enzyme recognizes double-stranded RNA (dsRNA) and catalyzes the adenosine-to-inosine (A-to-I) editing process. The binding of the sensor RNA to the target RNA and the formation of a dsRNA complex result in the recruitment of the ADAR enzyme to the dsRNA complex. The ADAR enzyme edits the adenosine in the UAG stop codon, converting it to inosine and thereby enabling translation of the downstream output protein (Fig. 1b). The primary requirement for designing the sensor sequence is thus to enable binding to the target RNA at a region containing a CCA motif. When the sensor and target RNAs form a dsRNA complex, this design aligns the UAG stop codon in the sensor with the CCA codon in the target, with a deliberate C-A mismatch at the center. This mismatch allows ADAR-mediated editing, which removes the STOP codon. To ensure equal expression levels of the reference and output proteins, the sensor RNA sequence is flanked on both sides by self-cleaving 2A peptides, resulting in the translation of a single transcript and the release of distinct protein products upon self-cleavage of the 2A peptide.

**Fig. 1.**
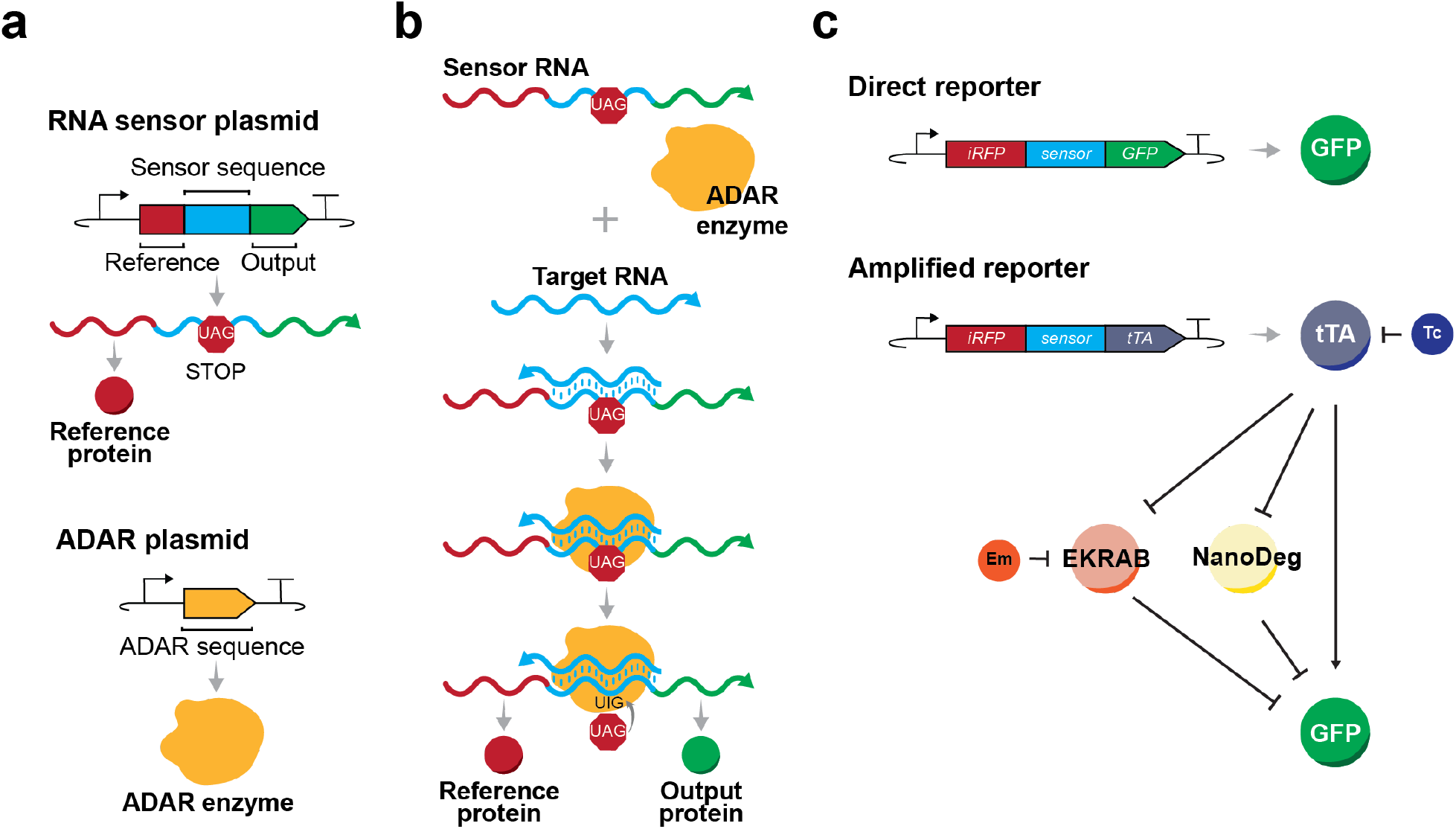
Design of an amplified RADAR reporter. **(a)** Schematic representation of the RADAR sensor. The RADAR sensor consists of two plasmids: one encoding the RNA sensor sequence (RNA sensor plasmid) and the other encoding the ADAR enzyme (ADAR plasmid). The RNA sensor plasmid encodes a tripartite sequence comprising a constitutively expressed reference gene, a sensor sequence complementary to the target RNA, and an output gene expressed upon detection of the target RNA. The sensor sequence is reverse complementary to the target RNA and contains a stop codon (UAG), leading to premature translation termination and preventing output translation. The ADAR plasmid encodes the ADAR enzyme, an adenosine deaminase that binds to double-stranded RNA and edits an adenosine into an inosine. **(b)** Mechanism of the RADAR sensor. Upon binding of the sensor sequence to the target RNA and the formation of a double-stranded RNA, the ADAR protein edits the double-stranded RNA to remove the stop codon, thereby leading to the translation of the output protein. **(c)** Comparison of a RADAR sensor with a direct output to a RADAR sensor with signal amplification. The RADAR sensor with direct output is based on a plasmid containing *iRFP* as a reference gene and *GFP* as an output gene. The RADAR sensor with signal amplification is based on a plasmid containing *iRFP* as a reference gene and *tTA* as an output gene. The RADAR sensor with tTA as output is co-expressed with negative transcriptional (EKRAB) and translational (NanoDeg) regulators of *GFP* under the control of tTA and *GFP* controlled by a hybrid promoter activated by tTA and repressed by EKRAB. tTA is regulated by tetracycline (Tc), and EKRAB is regulated by erythromycin (Em).

To assess strategies for optimizing the dynamic range and limit of detection of the RADAR sensor, we evaluated two sensor designs: one that produces GFP as output protein, and another that generates a transcription factor serving a master regulator in a signal amplification circuit, which is expected to improve both performance metrics (Fig. 1c).^28^ This signal amplifier platform is designed to enhance the dynamic resolution of the input by incorporating orthogonal regulatory elements that function at both transcriptional and post-translational levels to control output expression. Transcriptional regulation is expected to enhance the RADAR sensitivity by amplifying the output signal, while post-translational regulation is expected to refine the input dynamic resolution by overcoming limitations imposed by the intrinsic stability of the reporter protein. We implemented this strategy by engineering a RADAR sensor that produces a transcription factor—the tetracycline transactivator (tTA)—as its output. We placed the reporter gene (*GFP*) under the positive control of tTA. This design is expected to amplify the signal triggered by target RNA detection, thereby increasing the dynamic range of the output. To suppress background, leaky GFP expression in the absence of the target RNA, we incorporated an additional transcriptional control layer using the erythromycin-dependent transrepressor (EKRAB) placed under negative control of tTA. The *GFP* gene was placed under the control of a hybrid promoter responsive to both transcriptional regulators, activated by tTA and repressed by EKRAB. To further suppress GFP expression in the absence of the target and, critically, to accelerate the decay of GFP signal following a reduction in target RNA levels, we introduced a GFP-specific NanoDeg under negative control of tTA. The NanoDeg enables post-translational regulation via targeted proteasomal degradation, enhancing both the output dynamic range and the system’s responsiveness to changes in input. The use of small-molecule-dependent transcription factors—tTA (regulated by tetracycline, Tc) and EKRAB (regulated by erythromycin, Em)—allows tunable control of GFP expression. Adjusting Tc and Em concentrations modulates the levels of active tTA and EKRAB, respectively. Optimal Tc and Em concentrations are expected to minimize basal GFP expression in the absence of the target RNA and maximize the dynamic change in GFP fluorescence upon target RNA detection.

To evaluate the effect of the signal amplifier platform on the RADAR sensor performance, we compared the “direct reporter” RADAR sensor with GFP as output to the “amplified reporter” RADAR sensor that outputs tTA, which activates a downstream signal amplifier circuit leading to GFP expression (Fig. 1c). The direct reporter RADAR sensor plasmids were transfected into HEK293 cells; the amplified reporter RADAR sensor plasmids were transfected into a derivative HEK293 cell line (tTA-GFP). The tTA-GFP cell line was engineered to express (1) *GFP* under the control of a hybrid *CMV* promoter comprising a *7TO* operator for tTA activation and an *ETR* operator for EKRAB repression, and (2) a bicistronic construct encoding *NanoDeg* and *EKRAB* separated by an *IRES* and regulated by a *CMV* promoter and *TO* operator. This cell line was generated by lentiviral integration of the circuit components into HEK293 cells, followed by selection and clonal expansion.

### Integrating a signal amplifier with a RADAR sensor improves the RADAR sensor’s output dynamic range

To optimize the dynamic range of the amplified reporter RADAR sensor, we first systematically tested a range of Tc and Em concentrations to identify conditions that minimize GFP expression in the absence of target RNA and maximize the increase in GFP expression upon target RNA detection. We designed a RADAR sensor to detect the mouse *Bdnf* gene, which is not naturally expressed in HEK293 cells. To achieve controlled expression of the target RNA, we transfected tTA-GFP cells—containing the gene signal amplifier circuit—with a plasmid encoding mouse *Bdnf*, along with the amplified reporter RADAR sensor plasmid and the ADAR plasmid. Cells were transfected with the *Bdnf* RNA plasmid (Fig. 2a), and non-transfected cells were used as controls (Fig. 2b). A total of 30 Tc/Em combinations were tested, comprising five Em concentrations (0, 0.001, 0.01, 0.1, and 10 µg/mL) and six Tc concentrations (0, 0.0001, 0.001, 0.01, 0.1, and 10 µg/mL). GFP fluorescence was measured using flow cytometry (Fig. 2a-b and Supplementary Fig. 2). The highest fold change in GFP expression, a 12-fold increase, was observed in cells expressing *Bdnf* RNA treated with 0.01 µg/mL Tc and 10 µg/mL Em, indicating these small molecule concentrations most effectively enhanced the sensor dynamic range (Fig. 2c). Importantly, the optimal concentrations of Em and Tc depend on the specific target gene, particularly its basal expression level and the expected fold change.^28^ Therefore, small molecule concentrations must be individually optimized for each gene.

**Fig. 2.**
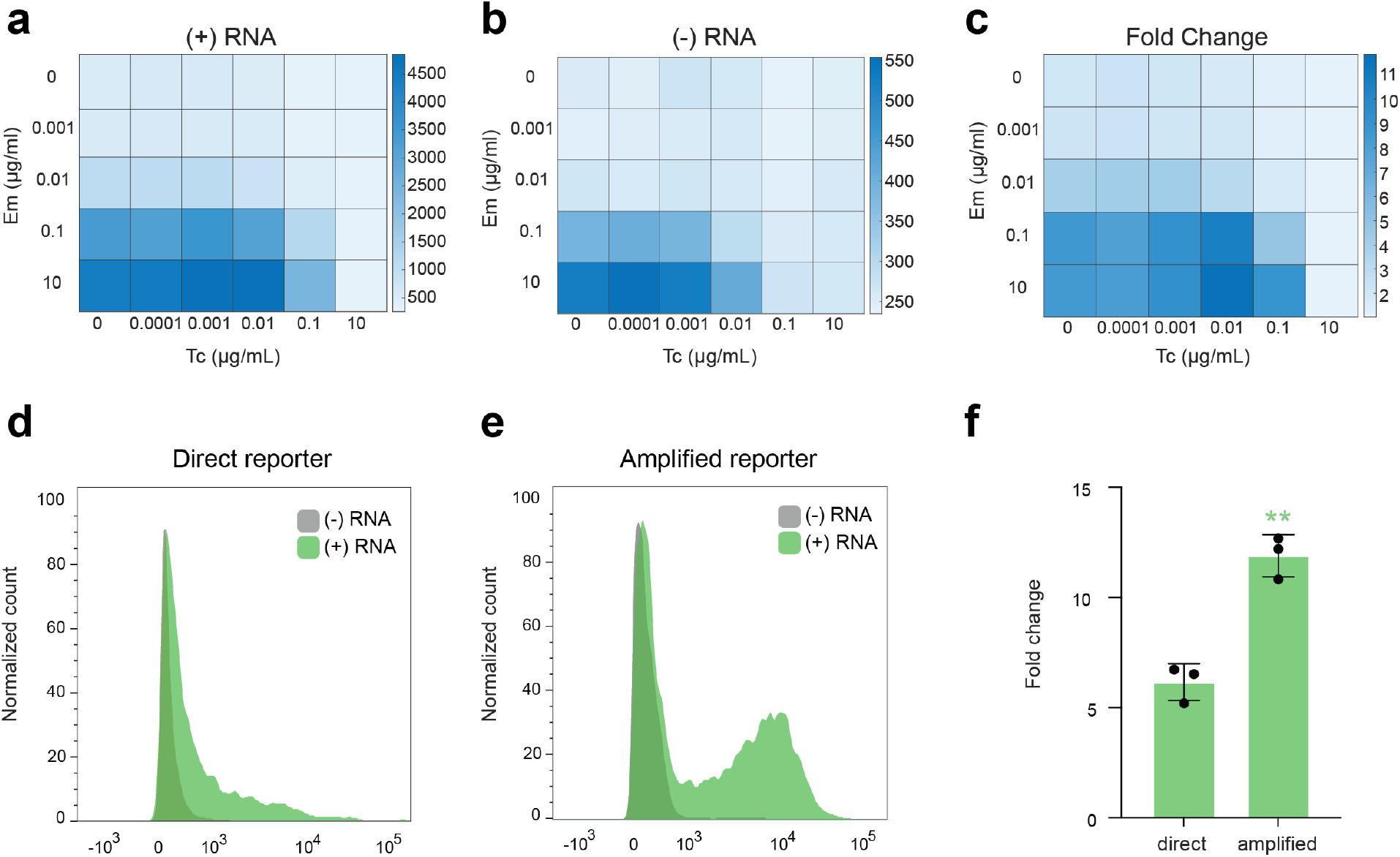
Characterization of the amplified RADAR reporter’s dynamic range. **(a-b)** Flow cytometry analysis of cells expressing the RADAR sensor coupled to the output signal amplifier circuit as a function of Tc and Em concentration, reported as GFP fluorescence measurements of tTA-GFP cells transfected with the *Bdnf* RADAR sensor and ADAR plasmids, and with **(a)** control (empty) plasmid or **(b)** *Bdnf* encoding plasmid. Data are reported as mean (n = 3). **(c)** Fold change of GFP fluorescence was obtained by normalizing the GFP fluorescence measurements of cells transfected to express *Bdnf* to that of cells lacking *Bdnf*. Data are reported as mean (n = 3). Biological replicates are reported in Supplementary Figure 2. **(d-e)** Representative histograms of flow cytometry analyses of GFP fluorescence of **(d)** HEK293 cells and **(e)** tTA-GFP cells transfected with the amplified *Bdnf* reporter and with *Bdnf* encoding plasmid (+) or with a control (empty) plasmid (-). Additional biological replicates are shown in Supplementary Figure 3. **(f)** Fold change of GFP expression of cells as in **(d-e)**. Fold change values were obtained by normalizing the GFP signal of cells transfected with the *Bdnf* encoding plasmid to that of cells transfected with the empty plasmid. Data are reported as mean (n = 3), (\*\**P* < 0.01).

To assess the impact of the signal amplifier on the dynamic range of the RADAR sensor, we compared the GFP output of the amplified reporter RADAR sensor to that of the direct reporter RADAR sensor. tTA-GFP cells were transfected as described above with a RADAR sensor targeting the *Bdnf* gene and producing tTA as output. Parental HEK293 cells were transfected with a RADAR sensor targeting the *Bdnf* gene and producing GFP as output. Both samples were also transfected with plasmids encoding the ADAR enzyme and the mouse *Bdnf* gene. Control cells lacking the *Bdnf* RNA were also included. Cells engineered with the amplified reporter were treated with optimized concentrations of Tc (0.01 µg/mL) and Em (10 µg/mL). GFP fluorescence was measured using flow cytometry. The direct reporter showed a 6-fold increase in GFP signal relative to controls, while the amplified reporter showed a 12-fold increase, demonstrating that the signal amplifier effectively doubles the output dynamic range (Fig. 2d–f and Supplementary Fig. 3).

### Integrating a signal amplifier with a RADAR sensor enhances sensitivity to the expression of native unfolded protein response (UPR) gene markers

To assess how signal amplification affects the limit of detection of RADAR sensors for endogenous RNA changes, we evaluated the expression of genes that are markers of the UPR, which can be induced chemically using the canonical UPR inducer tunicamycin.^28–30^ We selected four UPR-associated genes, namely *ERdj4, EIF4, BIP*, and *XBP1*, which are commonly used as markers of proteotoxic stress^28^ and are upregulated upon cell treatment with tunicamycin. In HEK293 cells, the basal transcript levels of these genes vary widely, with *ERdj4* measured at 7 nano transcripts per million (nTPM), *EIF4* at 179 nTPM, and *BIP* at 200 nTPM.^31^ For *XBP1*, UPR activation induces the formation of the spliced form of XBP1 mRNA (XBP1s). To establish a benchmark for evaluating the limit of detection of the RADAR sensors, we first quantified changes in the expression of the four genes upon treatment with tunicamycin. tTA-GFP cells were transfected with RADAR sensors targeting each UPR gene, using tTA as the reporter output. In parallel, parental HEK293 cells were transfected with the same sensors, but with GFP as the direct reporter. Both cell lines were also co-transfected with the ADAR-encoding plasmid. HEK293 and tTA-GFP cells were treated with tunicamycin (5 µg/mL) for 72 hours, and mRNA levels were quantified using RT-qPCR (Fig. 3a–b). Untreated cells were included as controls. The Ct values were normalized using measurements of *RNA18SN1* (18S RNA) and *ACTB* (Actin). Tunicamycin treatment led to a 1.9-, 2-, 4-, and 2.6-fold increase in ERdj4, EIF4, BIP, and XBP1s mRNA levels, respectively, in HEK293 cells. In tTA-GFP cells, corresponding increases were 2.2-, 2.2-, 3.8-, and 2-fold. These results confirm robust, tunicamycin-induced upregulation of UPR genes in both cell lines, establishing a reliable system for evaluating RADAR sensor sensitivity and detection limits for endogenous transcripts.

**Fig. 3.**
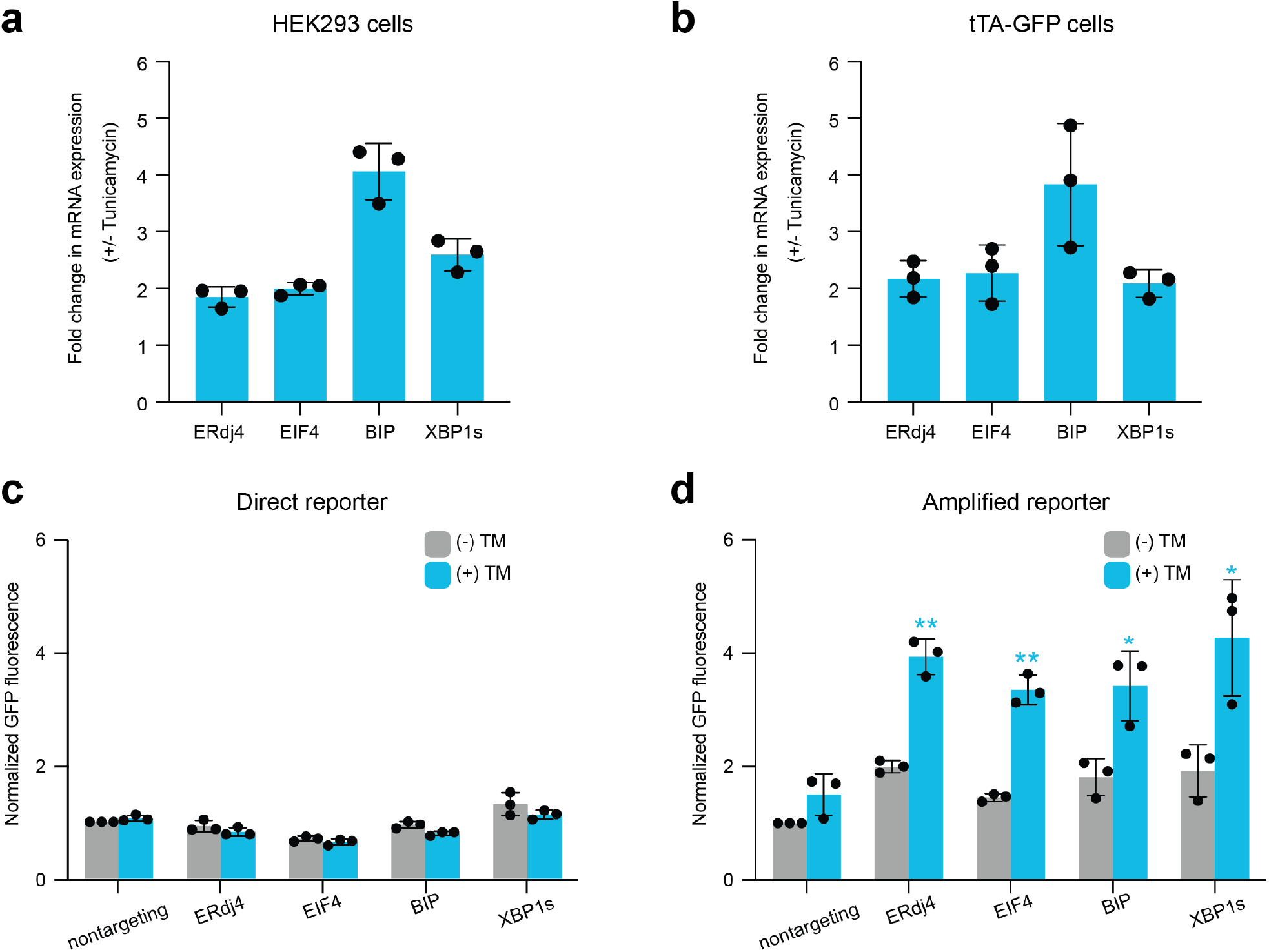
Detection of UPR-associated gene levels using direct and amplified RADAR reporters. **(a–b)** RT-qPCR analysis of *ERdj4, EIF4, BIP, and XBP1s* mRNA levels in **(a)** HEK293 cells transfected with direct reporters of *ERdj4, EIF4, BIP, and XBP1s* and **(b**) tTA-GFP cells transfected with amplified reporters of *ERdj4, EIF4, BIP, and XBP1s*. mRNA expression values were generated by normalizing the Ct values of the target genes to those of *RNA18SN1* (18S RNA) and *ACTB* (Actin) genes. Fold change values were obtained by normalizing mRNA levels of cells treated with tunicamycin (TM) (5 µg/mL, 72 h) to those of untreated cells. Data are reported as mean (n=3). **(c-d)** Flow cytometry analysis of **(c)** HEK293 cells transfected with the direct *ERdj4, EIF4, BIP*, and *XBP1s* reporters and **(d)** tTA-GFP cells transfected with amplified *ERdj4, EIF4, BIP*, and *XBP1s* reporters untreated (-) or treated with tunicamycin (5 μg/mL, 72 h) (+). GFP fluorescence values were normalized to the appropriate untreated nontargeting controls. Data are reported as mean (n=3), (\*\**P* < 0.01, \**P* < 0.05).

To assess the impact of signal amplification on the sensitivity and dynamic range of RADAR sensors for detecting endogenous UPR gene expression, we compared the amplified reporter RADAR sensors to the direct reporter sensors prepared as described above. A nontargeting RADAR sensor specific to *Bdnf* served as a negative control in both cell lines to normalize basal GFP fluorescence. A RADAR sensor lacking a stop codon was also used in both cell lines to assess any changes in GFP signal in response to tunicamycin treatment. Cells were treated with tunicamycin (5 µg/mL) for 72 hours or left untreated as controls. Cells expressing the amplified reporter sensors were also treated with the optimal doses of small-molecule inducers, specifically Em (1 µg/mL), which were determined through a systematic analysis of 30 Tc/Em combinations, as described above (data not shown). The resulting optimized small-molecule conditions, which do not require Tc, align with the inherently low basal transcript levels of UPR genes, consistent with the role of Tc in suppressing background activity under basal conditions. GFP fluorescence was measured by flow cytometry. The direct reporter sensors did not detect significant changes in target RNA levels upon UPR induction. In contrast, the amplified reporter sensors exhibited a significant GFP signal increase in response to tunicamycin treatment, 2- to 4-fold for *Erdj4*, 1.5- to 3.4-fold for *EIF4*, 1.8- to 3.4-fold for *BIP*, and 1.9- to 4.3-fold for *XBP1s* (Fig. 3c-d and Supplementary Fig. 4). Tunicamycin treatment did not lead to any change in signal in the control sensors lacking the STOP codon, demonstrating that tunicamycin treatment itself did not lead to a GFP signal increase (Supplementary Fig. 4). These findings indicate that signal amplification substantially improves RADAR sensor performance, enhancing both output dynamic range and sensitivity to changes in endogenous RNA expression, highlighting the utility of amplified reporter systems for monitoring native gene expression. Notably, this approach not only provides a reliable method for monitoring changes in transcriptional levels, but also for detecting splicing events such as IRE1-mediated XBP1s exon-exon junction.

### Integrating a signal amplifier with a RADAR sensor improves sensitivity to cobalt chloride-responsive gene expression

To further validate the effect of signal amplification on the limit of detection of RADAR sensors for endogenous RNA changes, we measured the expression of two hypoxia response marker genes, *HMOX1* and *BNIP3*. We induced hypoxia using the chemical hypoxia mimetic cobalt-chloride, which acts by stabilizing the hypoxia-inducible factor1 alpha (HIF-1α),^32–34^ leading to upregulation of these genes. In untreated HEK293 cells, the basal transcript levels of *HMOX1* and *BNIP3* are 8.7 nTPM and 46.6 nTPM, respectively.^31^

To assess the impact of signal amplification on the sensitivity and dynamic range of RADAR sensors, we compared the amplified reporter to the direct reporter in sensors designed to detect hypoxia-response genes. tTA-GFP cells were transfected with the RADAR sensors targeting the genes *HMOX1* and *BNIP3* and producing tTA as output. In parallel, parental HEK293 cells were transfected with RADAR sensors targeting the same genes, but designed to produce GFP as output. All samples were co-transfected with the ADAR-expressing plasmid. As before, we included a nontargeting RADAR sensor and a RADAR sensor lacking the STOP codon.

Cells were treated with cobalt chloride (200 µM) for 48 hours or left untreated as controls. Cells expressing the amplified reporter sensors were also treated with the optimal doses of small-molecule inducers (Em 1 µg/mL), determined through a systematic analysis of 30 Tc/Em combinations, as described above (data not shown). GFP fluorescence was measured by flow cytometry. The direct reporter sensors did not detect significant changes in target RNA levels upon UPR induction. In contrast, the amplified reporter sensors showed a significant increase in GFP signal upon tunicamycin treatment, with a 2.1- to 3.8-fold increase for *HMOX1* and 3.7- to 9-fold increase for *BNIP3* (Fig. 4a-b and Supplementary Fig. 5). Cells transfected with control sensors did not show significant changes in GFP signal, demonstrating that cobalt chloride treatment itself does not affect the RADAR output (Supplementary Fig. 5).

**Fig. 4.**
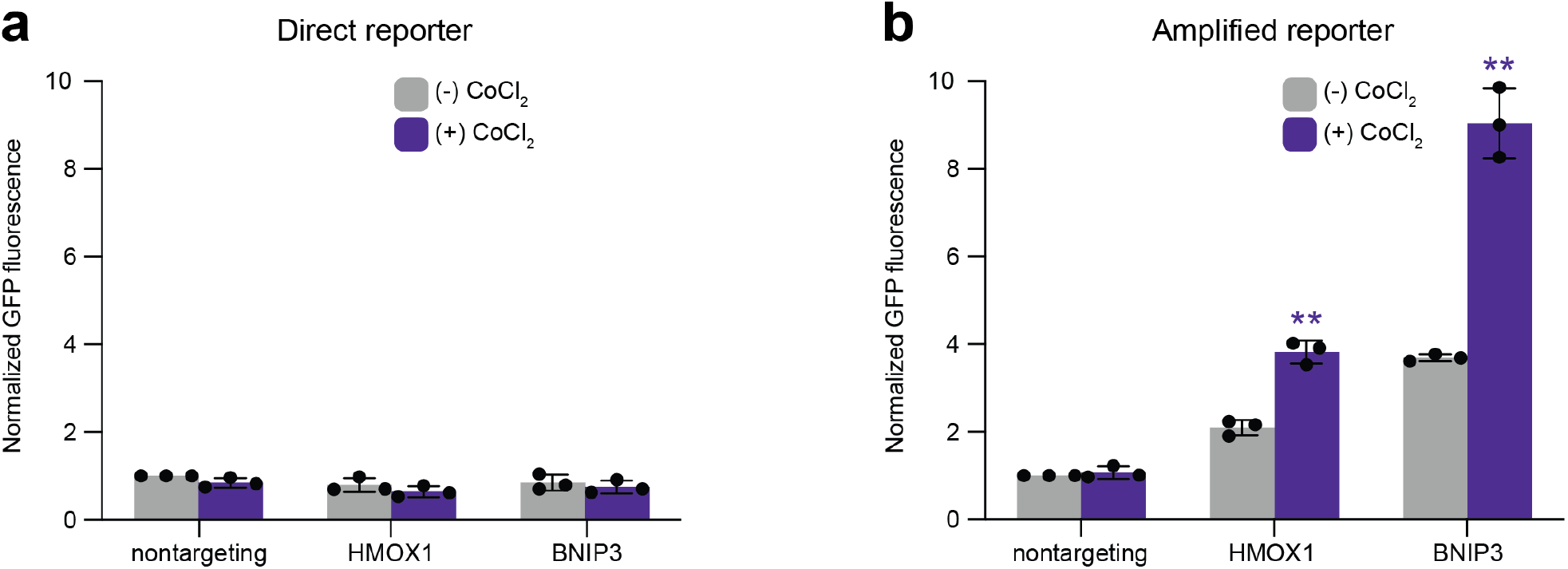
Detection of hypoxia response genes using direct and amplified RADAR reporters. **(a-b)** Flow cytometry analysis of **(a)** HEK293 cells transfected with direct *HMOX1* and *BNIP3* reporters and **(b)** tTA-GFP cells transfected with amplified *HMOX1* and *BNIP3* reporters, untreated (-) and treated with cobalt chloride (CoCl_2_) (200 μM, 48 h) (+). GFP fluorescence values were normalized to the appropriate untreated nontargeting controls. Data are reported as mean (n=3), (\*\**P* < 0.01).

Together with the earlier results demonstrating improved detection of UPR-related genes, these findings show that integrating the RADAR sensor with an orthogonal signal amplifier enhances the overall sensor’s dynamic range and sensitivity for monitoring endogenous gene expression, thereby validating the utility of this approach across various biological response pathways.

## Discussion

In this study, we present a strategy to significantly enhance the output dynamic range and sensitivity of RADAR sensors by integrating them with a genetic signal amplifier platform. This genetic circuit, which employs orthogonal regulatory elements at both transcriptional and post-translational levels, was designed to amplify the signal output of a gene reporter. To demonstrate its effectiveness, we compared a direct GFP reporter RADAR sensor with an amplified version that utilizes tTA to activate a downstream amplification circuit, resulting in enhanced GFP expression. Following a systematic optimization of the platform’s dynamic range by screening concentrations of key regulatory molecules, we confirmed that the signal amplifier significantly enhances the dynamic range of the RADAR sensors’ output. We further validated the system by designing RADAR sensors for a variety of endogenous genes, including those involved in the unfolded protein response (UPR) and hypoxia. Our results demonstrate that the signal amplification circuit markedly improves sensor performance, increasing both the dynamic range and sensitivity to native RNA fluctuations. These findings underscore the value of amplified reporter systems for precisely monitoring native gene expression.

We anticipate that our novel signal amplifier approach will complement other recent efforts to enhance the performance of RADAR. For example, similar to our work, previous studies have employed a binary vector approach, where a “reader” vector expresses tTA as an output protein and a separate “reporter” vector encodes GFP under control of a tetracycline-responsive element.^35,36^ These constructs, delivered using AAV vectors into neuronal cells, enable modular signal detection and reporting. Our platform, however, advances this concept by providing more precise control over leaky GFP expression and stronger amplification of the output signal. Additional strategies to boost RADAR performance, such as the co-expression of an ADAR2 payload targeted to the RNA edit site via MS2 RNA hairpin–coat protein interactions, could also be seamlessly integrated with our approach.^37^ Further enhancements can be achieved by optimizing sensor RNA design or by engineering RNA secondary structures that recruit endogenous RNAs, potentially improving both sensitivity and specificity^38^.

We anticipate that the approach presented herein, which integrates a signal amplifier into a sensor platform, will provide a broadly generalizable strategy to enhance the performance of other RNA-sensing technologies,^23,39^ as it can be easily implemented by swapping the sensor’s output with the master regulator, tTA. Importantly, it provides an alternative for enhancing sensor performance without requiring time-consuming biomolecular engineering of the sensor mechanisms themselves. Future work can focus on seamlessly integrating this platform with existing RNA sensors to improve their sensitivity and overall functionality, thereby validating its utility as a universal enhancement approach. Additionally, engineering compact genetic circuit cassettes that encode the signal amplifier platform enables co-delivery alongside RNA sensors, facilitating clinical translation. Finally, beyond cell-classifier applications, we anticipate the signal amplifier can be applied to boost therapeutic outputs in cell-based therapies^40–42^,where engineered cells serve as living therapeutics, thereby amplifying functional outcomes *in vivo*. In summary, integrating the signal amplifier platform offers a versatile and powerful framework for not only advancing RNA sensor technologies but also enhancing a broad range of therapeutic strategies.

## Methods

### Reagents

Tunicamycin (catalog no. T7765-5MG; Sigma-Aldrich) and erythromycin (catalog no. E5389-5G, Sigma-Aldrich) were dissolved in DMSO (catalog no. 472301; Sigma-Aldrich) to prepare a 10 mg/ml stock solution. Tetracycline (catalog no. T7660-5G; Sigma-Aldrich) was dissolved in water to prepare a 10 mg/ml stock solution. Cobalt chloride (catalog no. C8661-25G; Sigma-Aldrich) was dissolved in water to prepare a 12 mM stock.

### Plasmid assembly

All plasmids used in this study are listed in Table S1, with key sequences provided in Tables S2 and S3. Plasmid construction was performed using NEB Turbo *E. coli*. Addgene plasmids 191151 and 191141 served as backbones for constructing pJEC1614 and pJEC1615 by replacing the mCherry sequence with iRFP via Gibson Assembly, thereby reducing spectral overlap with GFP. Subsequently, pJEC1614 and pJEC1615 were used as templates to generate pJEC1616 and pJEC1617, in which GFP was replaced with a tTA sequence containing both a nuclear localization signal and a degradation tag. To create Golden Gate-compatible backbones for RADAR sensors, two BsaI sites were introduced into pJEC1615 and pJEC1617 using inverse PCR, yielding pJEC1618 and pJEC1619. Plasmids pJEC1620, pJEC1622, pJEC1624, pJEC1626, pJEC1628, and pJEC1630 were then assembled via Golden Gate cloning using the pJEC1618 backbone, BsaI, and sensor RNA inserts generated by annealing oligonucleotides (Millipore Sigma). Similarly, plasmids pJEC1621, pJEC1623, pJEC1625, pJEC1627, pJEC1629, and pJEC1631 were constructed using the pJEC1619 backbone and the same Golden Gate strategy with BsaI and annealed sensor RNA inserts. All plasmid constructs were verified using Sanger sequencing or full-plasmid sequencing.

### Sensor RNA design

The sensor RNA was generated using a Python script adapted from a previously published method^26^ that identifies candidate sequences for each gene. The script creates reverse-complementary sensor RNAs targeting all possible CCA motifs—where CCA is centrally located within a 99-bp window—and systematically mutates start codons, stop codons, and BsaI recognition sites to ensure compatibility with Golden Gate cloning. Sensor RNA sequences were selected based on two criteria: (1) a low number of homopolymer regions, defined as stretches of three or more identical nucleotides, and (2) a short maximum homopolymer length within each sequence. These criteria help reduce synthesis and cloning issues associated with repetitive sequences.

### Cell culture and transfections

HEK293 cells (catalog no. CRL-1573, ATCC) were cultured in DMEM/High glucose (catalog no. SH30243.01, Hyclone), supplemented with 10% fetal bovine serum (FBS, catalog no. 25-514H, GenClone) and 1% penicillin–streptomycin–glutamine (catalog no. SV30082.01, Hyclone), and maintained at 37 °C and 5% CO2. Cells were passaged using phosphate-buffered saline (catalog no. 17–516 F, Lonza) and trypsin (TrypLE Express, catalog no. 12605–036, GIBCO). The cell line tTA-GFP was generated in previous studies.

Transient transfections were conducted by seeding cells onto 6-well or 48-well plates. After 24 hours, upon reaching 70-80% confluency, transfections were performed using the JetPrime DNA transfection reagent (catalog no. 101000046; Polyplus) according to the manufacturer’s protocol. The medium was replaced 16 hours post-transfection. Cells were analyzed 48 hours post-transfection unless otherwise indicated.

### Flow cytometry analyses

Cells were analyzed using a FACSCanto II flow cytometer (BD Biosciences) and BD FACSDiva Software (v8.0.2). GFP fluorescence intensity was detected using a 488 nm laser and 530/30 nm emission filter. iRFP fluorescence intensity was detected using a 635 nm laser and 780/60 nm emission filter. Cells were first gated on a linear forward scatter (FSC-A) area versus linear side scatter (SSC-A) plot to identify the main cell population. To exclude cellular debris and other non-cellular events, the bottom 20% of events of both FSC-A and SSC-A were removed. Doublets and cell aggregates were then excluded by gating on the forward scatter area (FSC-A) versus forward scatter height (FSC-H) plot. After gating for singlets, the GFP fluorescence of the gated iRFP-positive population was analyzed. A minimum of 10,000 singlet cells or 2,000 iRFP-positive events were recorded in each sample. Data analysis was performed using FlowJo (v10.10.0) and Microsoft Excel (v25040241).

### RT-qPCR experiments

RNA was extracted using the RNeasy Plus Mini Kit (catalog no. 74134, Qiagen), and cDNA was synthesized using the qScript cDNA SuperMix (catalog no. 95048-100, Quanta Biosciences) following the manufacturer’s protocol. RT-qPCR reactions were performed using the PerfeCTa SYBR Green FastMix (catalog no. 95072-012, Quanta Biosciences) in a CFX96 Real-Time PCR Detection System (Bio-Rad) using appropriate primers (Table S2). The housekeeping genes *RNA18SN1* and *GAPDH* were used as reference points to normalize the data.

### Statistical analyses

All data are reported as mean, and error bars represent the standard error of at least three independent experiments. For each replicate, fluorescence was measured using flow cytometry, and the mean fluorescence intensity of the population was used for analysis. Heatmap representations of data were created using MATLAB software (MathWorks) using the mean of three biological replicates. The statistical significance of experiments was calculated using a two-tailed Student’s t-test. No statistical method was used to predetermine the sample size. No data were excluded from the analyses, the experiments were not randomized, and the investigators were not blinded to allocation during the experiments and outcome assessment.

## Abbreviations

(RADAR): RNA sensing using Adenosine Deaminases Acting on RNA
(UPR): Unfolded protein response
(HEK293 cells): Human embryonic kidney 293 cells
(tTA): tetracycline transactivator
(EKRAB): erythromycin-dependent transrepressor
(Tc): tetracycline
(Em): erythromycin
(TM): tunicamycin

## Author Information

### Author Contributions

H.L, J.C., and L.S. conceived the study. H.L. performed all experiments and analyzed the data. All authors wrote the manuscript. H.L, J.C., and L.S. wrote the manuscript.

### Supporting Information

Additional experimental details and measurements, and plasmids, DNA sequences, and primers (PDF)

Supporting data (XLSX)

## Acknowledgement

This project was supported by National Science Foundation Award 2128370 (J.C., L.S.), the Robert J. Kleberg, Jr. and Helen C. Kleberg Foundation award A23-0202-004 (J.C.), and by the National Institutes of Health Award EB030030 (L.S.).

